# Material perception relies on context-dependent active sensing strategies

**DOI:** 10.64898/2025.12.19.695393

**Authors:** Ryu Nomachi, Hideki Tamura, Kevin A. Helgeland, Takuma Morimoto, Shigeki Nakauchi, Tetsuto Minami

**Author notes:** Correspondence concerning this article should be addressed to Hideki Tamura, 1-1 Hibarigaoka, Tempaku-cho, Toyohashi, Aichi, JAPAN, 441-8580. These authors contributed equally to this work (Ryu Nomachi, Hideki Tamura). The authors made the following contributions. Ryu Nomachi: conceptualization, data curation, formal analysis, investigation, methodology, software, validation, visualization, writing - original draft, writing - review & editing; Hideki Tamura: conceptualization, data curation, formal analysis, funding acquisition, investigation, methodology, project administration, resources, software, supervision, validation, visualization, writing - original draft, writing - review & editing; Kevin A. Helgeland: data curation, formal analysis, investigation, methodology, software, validation, visualization, writing - original draft, writing - review & editing; Takuma Morimoto: conceptualization, investigation, supervision, validation, writing - review & editing; Shigeki Nakauchi: funding acquisition, resources, supervision, validation, writing - review & editing; Tetsuto Minami: funding acquisition, resources, supervision, validation, writing - review & editing.

## Abstract

Material perception is typically studied under passive and highly constrained viewing conditions, thus leaving it unclear whether humans rely on active sampling strategies to resolve perceptual ambiguities. In everyday vision, however, observers naturally move their heads and manipulate objects with their hands to obtain informative cues, thus raising the question of how such exploratory actions contribute to material recognition. We used immersive virtual reality to examine whether viewpoint changes and object manipulations support the discrimination of visually challenging materials, specifically metal versus glass, whose appearances depend strongly on the associated illumination and viewing geometry. On the basis of three experiments, we revealed that observers systematically increased their exploration tendencies when the identity of a material was ambiguous and that this increased movement was associated with higher discrimination accuracy. By independently manipulating head- and hand-based motions, we identified the context-dependent contributions of each modality, revealing that viewpoint changes dominated when multiple objects could be compared simultaneously, whereas object manipulation was more effective when only a single target was available. Moreover, the participants differed in terms of the efficiency of their sampling strategies, and those who employed more informative exploration patterns exhibited both higher accuracy and greater performance gains across different trials. These results demonstrate that material recognition depends on flexible, context-dependent active sensing rather than passive evaluations of static images. Our findings introduce a new framework, according to which material perception emerges from an adaptive perception–action loop, thus highlighting the functional role of exploratory behaviors in complex visual environments.

**Significance Statement:** The task of material perception in natural environments depends not only on the optical properties of surfaces but also on the actions that observers take to reveal informative visual cues. However, most laboratory studies rely on passive viewing, and how humans actively regulate their own movements to reduce perceptual uncertainty remains unknown. On the basis of three virtual-reality experiments, we showed that observers flexibly perform head- and hand-based explorations to enhance the discriminability of visually ambiguous materials. These exploratory strategies were strongly context-dependent and exhibited systematic individual differences that predicted learning-related improvements. Our findings demonstrate that material perception emerges from an adaptive perception–action loop rather than static image analyses, providing a framework for understanding how humans actively acquire diagnostic sensory information in complex environments.

## Introduction

Material perception allows humans to identify what objects are made of—such as metal, glass, plastic, or fabric—based on visual information. A large body of work has established how mid-level visual processes recover surface properties from the interplay of shading, specular reflections, and three-dimensional shapes (1–5). These studies, which are grounded in image-computable analyses and computational models (6–13), have shown that many of the cues supporting material perception are inherently dynamic, changing systematically with illumination conditions (14, 15), viewpoint (16–18), and object poses (19–24). Such dynamic transformations—including specular flow patterns, highlighted motions, and refractive distortions—often provide the diagnostic information that is necessary to distinguish materials in static images that are otherwise ambiguous.

Despite this progress, nearly all empirical and theoretical research on material perception has relied on passive or experimenter-controlled viewing conditions. Observers typically remain stationary while objects are rotated for them, or they view brief animations in which motions are predetermined. These paradigms successfully reveal which optical cues can support material recognition, but they leave open a foundational question: How do humans actively sample visual information when they are free to select their own movements? In natural vision, observers rarely experience objects under fixed or externally controlled conditions. Instead, they continuously move their heads and hands to obtain informative viewpoints (Figure 1A). Such behaviors—approaching an object, rotating it, or peering at it from above or below—fundamentally change the retinal input. However, we know little about how people choose among these candidate actions, how they deploy them under perceptual uncertainty, or how different exploratory modalities contribute to material judgments. As a result, we currently lack a principled account of active material perceptions, although most real-world material judgments depend on self-generated viewpoints and pose changes.

**Figure 1.**
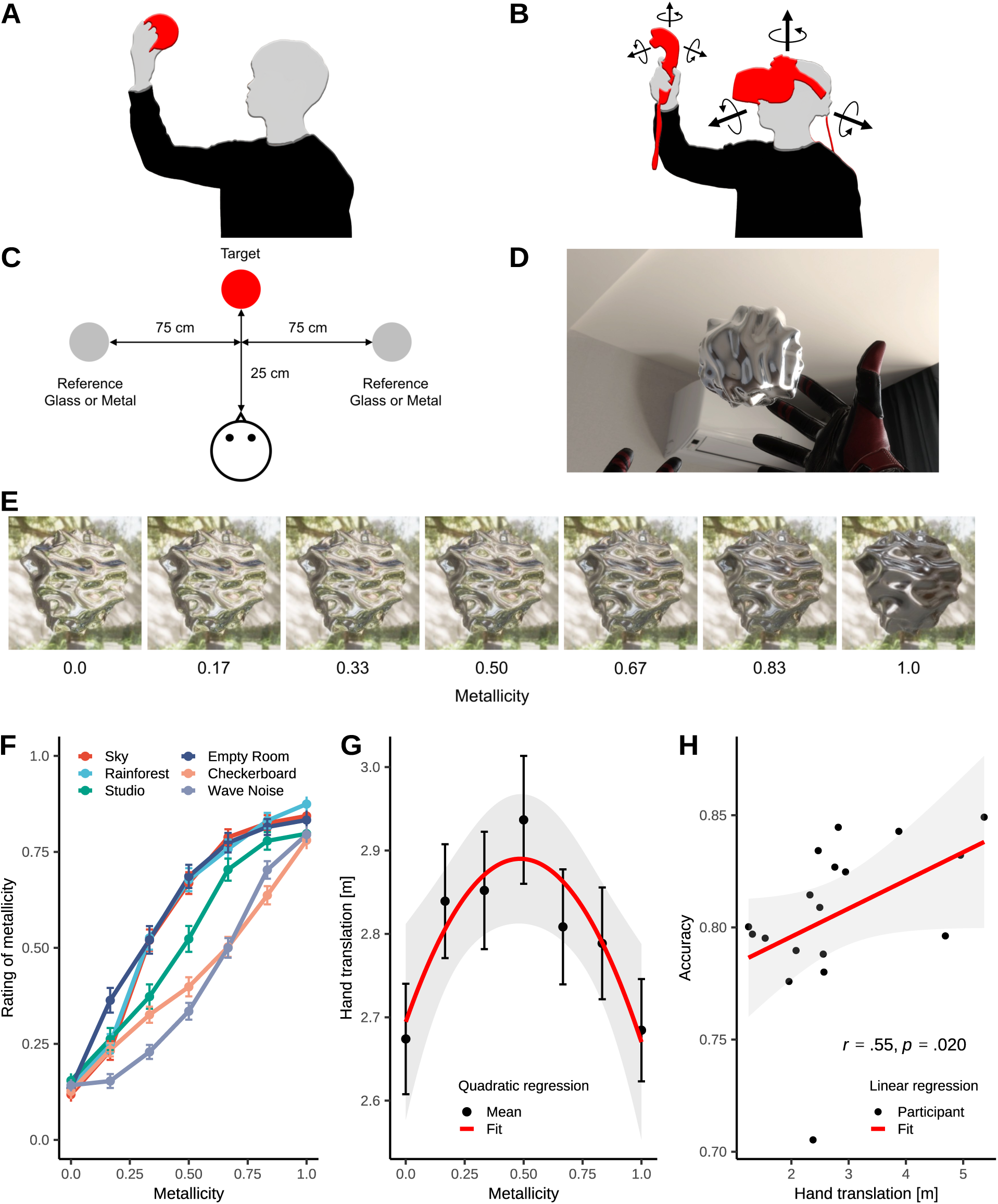
Experimental setup and results of Experiment 1. (A) Conceptual illustration of the study, depicting an observer freely exploring objects in a naturalistic environment. (B) Virtual-reality implementation of the scenario shown in (A). The head and hand positions and orientations of the participants were continuously recorded from a head-mounted display (HMD) and handheld controllers, respectively. (C) Schematic diagram of the experimental setup and the spatial arrangement of the stimuli and participants involved in Experiment 1. (D) Example of a participant’s view in Experiment 1, as rendered within the virtual environment. (E) Examples of the experimental stimuli, illustrating a continuum from a glass-like appearance (metallicity = 0.0) to a metal-like appearance (metallicity = 1.0). (F) Relationship between physical metallicity and the subjective perceived metallic appearance ratings. Line colors indicate different illumination environments. The error bars denote the standard errors. (G) Relationship between physical metallicity and the amount of hand translation. The error bars indicate the standard errors, and the shaded regions represent 95% confidence intervals. (H) Relationship between the amount of hand translation and the resulting discrimination accuracy.

Recent work (25) has suggested that allowing observers to actively manipulate objects can improve their perceptual decisions, particularly when making glossiness judgments that depend on specular structure information. However, these studies have not resolved why and when an active exploration becomes beneficial, whether its effectiveness depends on the structure of the viewing environment, or how different actions—such as viewpoint changes versus object manipulation—contribute to resolving material ambiguities. More broadly, perception–action coupling theories propose that exploratory movements are not merely outputs of the motor system but rather integral components of perceptual inference (26). Whether such principles apply to material perception remains unknown.

Therefore, the present study addresses this gap by introducing a virtual-reality (VR) framework that allows observers to freely move both their viewpoints and the objects that they are inspecting (Figure 1B). This approach enables fine-grained, frame-by-frame measurements of head and hand movements while preserving ecological validity. Utilizing this framework, we asked three fundamental questions. 1. Do observers strategically regulate their movements to resolve material ambiguities? 2. How do viewpoint changes (head motions) and object manipulations (hand motions) differentially contribute to the discrimination process? 3. Do individuals exhibit distinct exploratory strategies, and do these strategies predict learning effects and performance?

Across three experiments, participants judged whether visually challenging objects were metal or glass because the appearances of these materials are strongly modulated by environmental illumination condition (8, 14, 23, 24, 27, 28). Metal surfaces produce mirror-like reflections of the surrounding scene (29–33), whereas glass transmits and refracts light, creating distorted images of the background (34–40). These optical properties make the metal–glass distinction a particularly challenging material-recognition problem, especially when observers actively explore an object in relation to its environment. Furthermore, by manipulating whether head or hand motions produced visual consequences, we dissociated the contributions of each modality. We quantified exploratory behaviors using translation and rotation metrics, and extracted individual strategy profiles using principal component analysis (PCA) and Gaussian mixture model clustering. This allowed us to link how participants moved to how well they perceived objects, revealing the strategic dimensions of active sensing.

Our findings reveal that active explorations are not uniformly beneficial, but instead depends critically on the availability of informative dynamic cues. Viewpoint changes enhanced the discrimination capabilities of the participants when comparative cues were present, whereas object manipulation was more effective when the viewpoint cues were limited. Crucially, the number of exploratory movements increased selectively under conditions of high perceptual uncertainty, indicating that the observers choose actions that reduced the degree of ambiguity rather than producing nonspecific motions. Moreover, the individuals exhibited striking differences in terms of their exploratory strategies, and those who adopted more efficient sampling behaviors showed greater learning effects across different trials. Together, these results show that material perception emerges from a flexible, context-dependent perception–action loop, challenging the static-image assumptions that dominate the current theories. By integrating VR-based behavioral measurements with quantitative exploratory action modeling, our study provides a new framework for understanding how humans actively construct the visual information that is needed for recognizing materials in complex environments.

## Results

### Active Exploration Contributes to Material Discrimination

In Experiment 1, the target object appeared directly in front of the observer, with the reference objects, one with metallicity = 0.0 (glass) and the other with metallicity = 1.0 (metal), positioned 90° to the left and right (Figure 1C). By pressing the trigger on the controller in one hand, the participants could virtually “grasp” the target and move it freely in space. Under two object shapes (Figure S1A) and six illumination environments (Figure S1B), participants judged how similar the target appeared to each reference material using a continuous slider (the participants’ point of view is shown in Figure 1D). Metal and glass were chosen because their appearances depend strongly on illumination and viewing geometry conditions. Metals primarily convey their identity through specular reflections of the surrounding environment, whereas glass reveals refractive and transmitted light patterns. These properties make the metal–glass distinction a demanding perceptual judgment task, providing a suitable test case for examining how active explorations support material perceptions (example stimuli are presented in Figure 1E).

An analysis of the rating data (Figure 1F) revealed that higher physical metal contents led to higher perceived metallicity levels (*F* (1, 17) = 139.56, *p < .*001, *η*^2^ = 0.891), and that the judgments differed across illumination environments (*F* (5, 2977) = 18.13, *p < .*001, *η*^2^ = 0.03). We also observed a significant interaction between the material and illumination (*F* (5, 2977) = 4.25, *p < .*001, *η*^2^ = 0.007), indicating that the degree to which an object appeared metallic—or glass-like—depended strongly on the surrounding light field. Specifically, objects tended to be perceived as more metallic under natural illumination, whereas the same objects were more likely to be perceived as glass-like under artificial illumination. These results confirmed that, even under active viewing, observers could reliably discriminate between materials in a manner consistent with that observed in previous passive viewing studies.

Turning to the participants’ exploratory behaviors, we found that the amount of object translation exhibited a clear inverted U-shaped relationship with the physical material along the metal–glass continuum (Figure 1G): the degree of translation was greatest at metallicity = 0.50, where the stimulus was most ambiguous (*β* = −0.83, *p* = 0.002). This relationship was also observed within each illumination condition individually. These results suggest that the observers increased their exploratory movements when diagnostic cues were more difficult to obtain. Similarly, participants who translated the object more extensively achieved higher overall discrimination accuracy, defined as the normalized agreement between each participant’s rating and the true metallicity of the stimulus, yielding a positive correlation between exploration and performance (Figure 1H; *r* = 0.55, *p* = 0.020). Together, these findings indicate that active viewing affords observers the freedom to deploy sampling strategies, and that effective explorations directly contributes to improved material recognition results.

### Dissociating the Contributions of Viewpoint Motions and Object Manipulations

The exploratory behavior observed in Experiment 1 combined two sources of movement: viewpoint changes produced by head motions and object manipulations produced by hand motions. This made it unclear which modality-specific cues were primarily responsible for the material discrimination improvement observed in the results. Previous work published by Lin and colleagues (25) demonstrated that active explorations can benefit material judgments, and that observers generally conduct more extensive explorations during gloss judgments than during lightness judgments. However, the specific movements that drive this benefit—and the contexts in which they matter—remain unresolved.

To address this gap, Experiment 2 dissociated the head- and hand-based contributions by selectively enabling or disabling the motion signals derived from the HMD and controller. Utilizing a setup similar to that used in Experiment 1 but with the reference objects positioned on both sides of the target (Figure 2A), we tested all combinations of head motion tracking (on/off) and controller tracking (on/off), yielding four conditions that isolated the effects of viewpoint motions and object manipulations (Figure 2B). The participants performed a 2AFC task to determine whether the target appeared more similar to the left or right reference material (metal or glass) while varying the viewpoint, the object, both, or neither. The illumination conditions included two natural light fields and two artificial fields to examine whether the usefulness of each exploratory modality depended on the environmental context.

**Figure 2.**
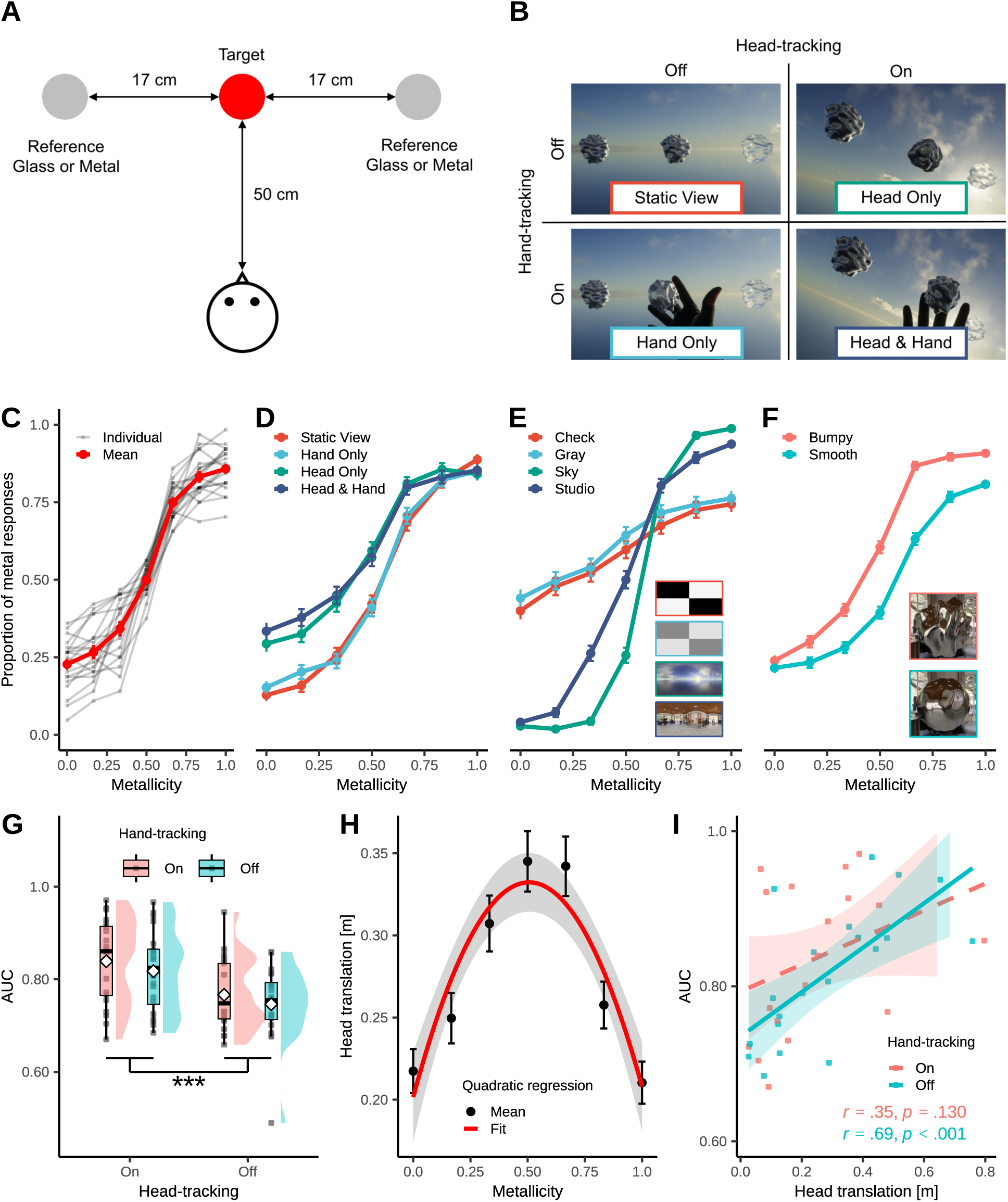
Results of Experiment 2. (A) Schematic illustration of the experimental setup and the spatial arrangement of the stimuli and participants involved in Experiment 2. (B) Experimental conditions and example views from the participants’ perspective in the virtual environment. (C) Relationship between physical metallicity and the subjective metal responses. The red line indicates the group mean, and the light gray lines represent individual participants’ data. (D–F) Same data as those shown in (C), replotted separately by experimental condition, illumination, and object shape. (G) Discrimination performance indexed by the area under the ROC curve (AUC) across different experimental conditions. White diamonds denote group means, and gray points indicate individual participants. (H) Relationship between physical metallicity and the amount of head translation. The formatting is identical to that utilized in Figure 1G. (I) Relationship between the amount of head translation and the resulting discrimination performance (AUC). Asterisks indicate significant differences: ∗ ∗ ∗*p < .*001.

An analysis of the choice data (Figure 2C) revealed that a higher physical metal content again led to higher metal-response rates, replicating the results of Experiment 1. When the results were broken down by viewing condition, the participants were more likely to judge the stimuli as metallic when head motions were available (Figure 2D). The illumination level also modulated the responses: artificial light fields yielded more ambiguous judgments than natural fields did (Figure 2E). In addition, objects with deeper surface reliefs were more often judged as metallic (Figure 2F). Together, these results indicated that the material judgments depended strongly on the observer movements, illumination conditions, and shapes.

We subsequently examined the discrimination accuracy achieved across the four viewing conditions (Figure 2G). A significant main effect of head motion was observed (ART ANOVA: *F* (1, 76) = 13.83*, p < .*001), whereas the main effect of hand motion was not significant (*F* (1, 76) = 0.70*, p* = 0.406), and no interaction was detected (*F* (1, 76) = 0.10*, p* = 0.756). Thus, in this context—where participants could observe the target and reference stimuli simultaneously—the cues obtained through viewpoint changes played a dominant role. Similar to Experiment 1, the numbers of both head and hand translations peaked when the physical material was most ambiguous, indicating increased exploratory behavior under uncertainty (*β* = −0.51, *p < .*001 for head translation; Figure 2H). Crucially, more movements did not uniformly improve the resulting performance. A strong positive correlation was found between the head translation distance and discrimination accuracy only when hand motions were disabled (*r* = 0.69, *p < .*001; Figure 2I). When hand motions were available, this correlation weakened (*r* = 0.35, *p* = 0.128; Figure 2I). A similar pattern emerged for head rotations (no-hand condition: *r* = 0.72, *p < .*001; hand condition: *r* = 0.26, *p* = 0.271). These results indicate that, in Experiment 2, viewpoint motion served as the primary exploratory strategy because it provided simultaneous access to cues from both the target and reference objects—an advantage that was especially pronounced when object manipulations offered no additional information.

### When Viewpoint Changes Provide Limited Information, Observers Rely on Object Manipulation Instead

What strategies do observers adopt when viewpoint changes fail to provide sufficient diagnostic information? To address this question, in Experiment 3, the reference objects used in Experiment 2 were removed, leaving only the target object in view (Figure 3A). The participants performed a binary classification task to judge whether the target appeared metallic or glass-like while head-motion tracking and hand-motion tracking were independently enabled or disabled, as in Experiment 2 (Figure 3B). In this configuration, viewpoint motions could no longer yield comparative cues from the reference objects, reducing the relative utility of head-based explorations. If the participants continued to rely on viewpoint changes despite this limitation, it would imply that head motion constitutes a fundamentally important modality for active material perception tasks. Conversely, if they increased the amount of object manipulation and achieved higher discrimination performance in conditions where hand motions were available, this would indicate that observers flexibly shift their exploratory strategy depending on which cues are most informative in the given context.

**Figure 3.**
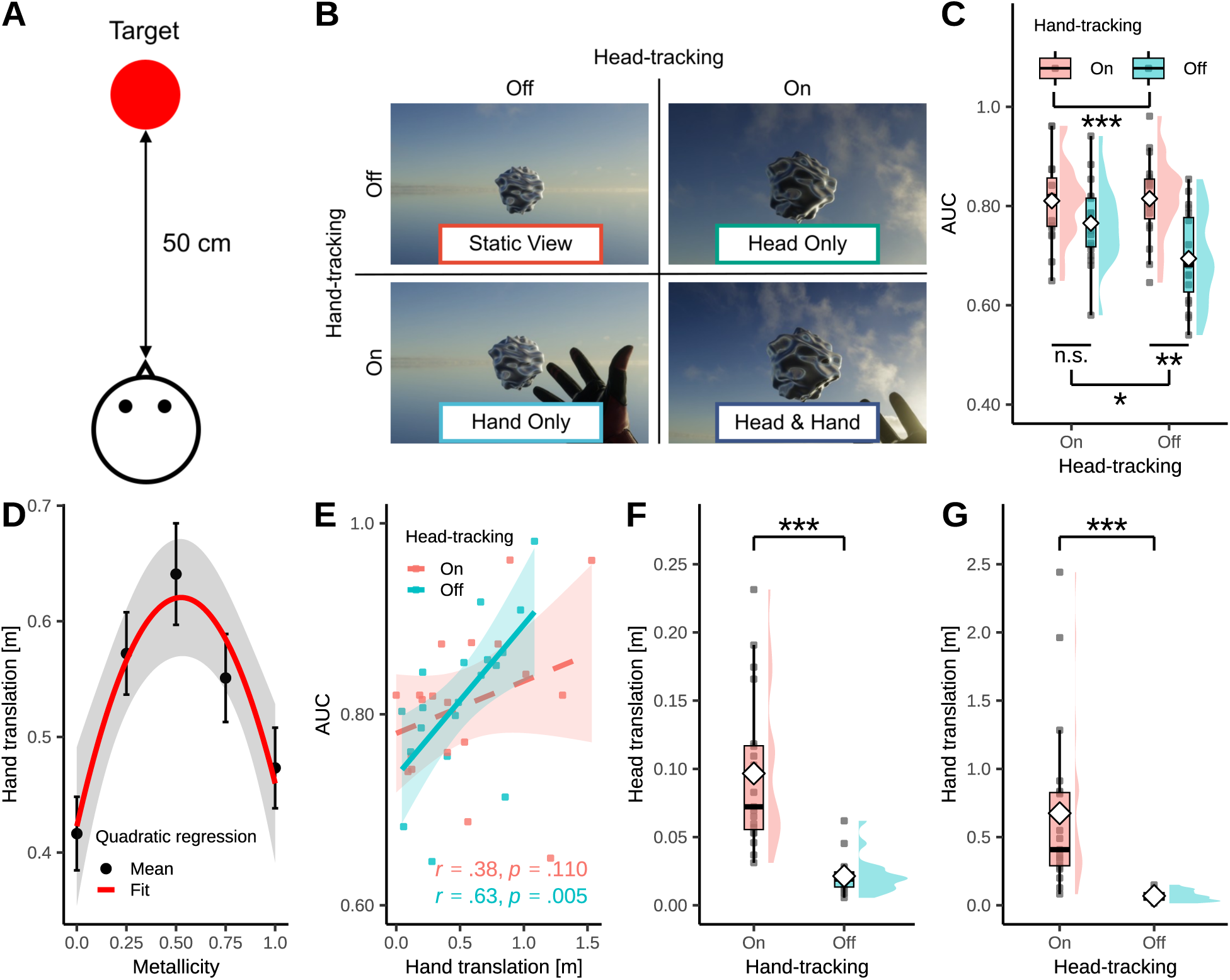
Results of Experiment 3. (A) Schematic illustration of the experimental setup in Experiment 3. (B) Experimental conditions and example participant-view images in Experiment 3. (C) Relationships between the experimental conditions and the resulting discrimination accuracy (AUC). (D) Relationship between the degree of metallicity and the amount of hand translation. (E) Relationship between the amount of hand translation and the AUC. Panels (C–E) follow the same format as employed Figures 2G, 2H, and 2I. (F) Head translation in the head-tracking-off condition (i.e., “empty” head movements). (G) Hand translation in the hand-tracking-off condition (i.e., “empty” hand movements). Notably, the identical test statistics in (F) and (G) were the result of the fact that the Wilcoxon signed-rank tests were applied to two measures derived from the same paired observations, resulting in the same rank ordering across participants. Asterisks indicate significant differences: ∗*p < .*05, ∗ ∗ *p < .*01., ∗ ∗ ∗*p < .*001.

We observed similar results in that the experimental conditions, illuminations, and object shapes affected the metal responses even when only the target object was presented (see Figures S1C and S1D for the illuminations and stimuli, respectively; see Figures S1E-S1H for the results). An analysis of the achieved discrimination accuracy (Figure 3C) revealed not only a significant main effect for head motion (*F* (1, 18) = 6.92, *p* = .017, *η*^2^ = 0.278) but also a significant main effect for hand motion (*F* (1, 18) = 18.19, *p < .*001, *η*^2^ = 0.503), as well as a significant interaction between the two factors (*F* (1, 18) = 5.53, *p* = .030, *η*^2^ = 0.235). Multiple comparisons revealed that hand motion improved the resulting discrimination accuracy only when head motion was disabled (*t*(18) = 4.71, 95% CI [0.04, 0.20], *p* = 0.001, *d* = 1.08), whereas no such benefit was observed when head motion was enabled (*t*(18) = 1.81, 95% CI [-0.03, 0.12], *p* = 0.519, *d* = 0.42). Thus, in contrast with Experiment 2, hand-based object manipulation contributed to the performance achieved in Experiment 3 only under conditions in which viewpoint-based cues were unavailable. In support of this interpretation, the object translation distance increased when the physical material was most ambiguous, indicating greater reliance on manipulation-based explorations (*β* = −0.72, *p < .*001; Figure 3D). Moreover, participants who moved the object more extensively achieved higher discrimination accuracy, but only when hand motion was available and head motion was disabled (*r* = 0.63, *p* = 0.004; Figure 3E). This relationship was absent when both head and hand motions were enabled (*r* = 0.38, *p* = 0.112; Figure 3E). These findings indicate that observers preferentially rely on object manipulations when they provide sufficiently diagnostic information, demonstrating flexible, context-dependent sampling strategies.

Curiously, when tracking for one modality was enabled, the number of movements in the other modality increased even when its tracking process was disabled (head translation: *V* = 190, *Z* = 4.62, *p < .*001, *r* = 0.75 in Figure 3F; hand translation: *V* = 190, *Z* = 4.62, *p < .*001, *r* = 0.75 in Figure 3G). In other words, under conditions where informative cues were limited, participants produced “empty” head or hand motions—movements that did not affect the visual stimulus—suggesting an effort to obtain missing information even when such actions were not visually effective.

### Individual Differences in Exploratory Strategies Predict Discrimination Performance

The results thus far have indicated that the observers flexibly switched between head-based and hand-based sampling strategies depending on the viewing context. We next asked how much these exploratory behaviors varied across individuals and which specific movement patterns were most conducive to achieving accurate material discrimination. To quantify these differences, we categorized each participant’s viewing behavior in terms of four components: “pulling the object closer,” “approaching the object,” “viewing the object from below,” and “viewing the object from above” (see Figure S2A for Experiment 2 and Figure S2B for Experiment 3). By reference to these behavioral metrics, we classified the participants according to the exploratory strategies they employed during the task.

An analysis of Experiment 2 conducted under conditions in which head-based exploration was most informative (Figure 2G) revealed marked individual exploratory strategy differences. A PCA followed by Gaussian mixture model (GMM) clustering (Figures 4A and 4B) revealed that the participants varied primarily along two dimensions: their overall observation times (PC1; 59.8% of the explained variance) and the relative balance between head- and hand-based explorations (PC2; 20.6% of the explained variance). On the basis of these dimensions, the participants were classified into three distinct clusters. The evolution trends observed for the discrimination accuracy across four successive trial blocks for each cluster are shown in Figure 4C. A two-way mixed-effects analysis with block and cluster factors revealed a significant main effect for clusters (*F* (2, 17) = 7.28, *p* = 0.005, 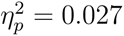), but no main effect for blocks (*F* (1, 57) = 1.58, *p* = 0.214, *η*^2^_*p*_ = 0.461) and no block × cluster interaction (*F* (2, 57) = 0.56, *p* = 0.577, 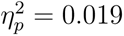). A visual inspection of the data (Figure 4C) further revealed that Cluster 1, which exhibited the strongest learning-related improvement (i.e., the greatest AUC increase from the first block to the final block), was characterized by a moderate exploration time and a pronounced tendency to approach the stimulus by moving the head toward it. This pattern indicates that under the conditions of Experiment 2, viewpoint-based strategies involving an active approach for addressing the object were particularly effective for material discrimination purposes. In contrast, Cluster 2 had the lowest discrimination accuracy. Although the overall exploration time was short, the participants in this cluster exhibited no clear preference for either approaching the object with their heads or pulling it closer to their hands, indicating low strategic consistency. This mixed and inefficient exploration pattern likely contributed to their reduced performance.

**Figure 4.**
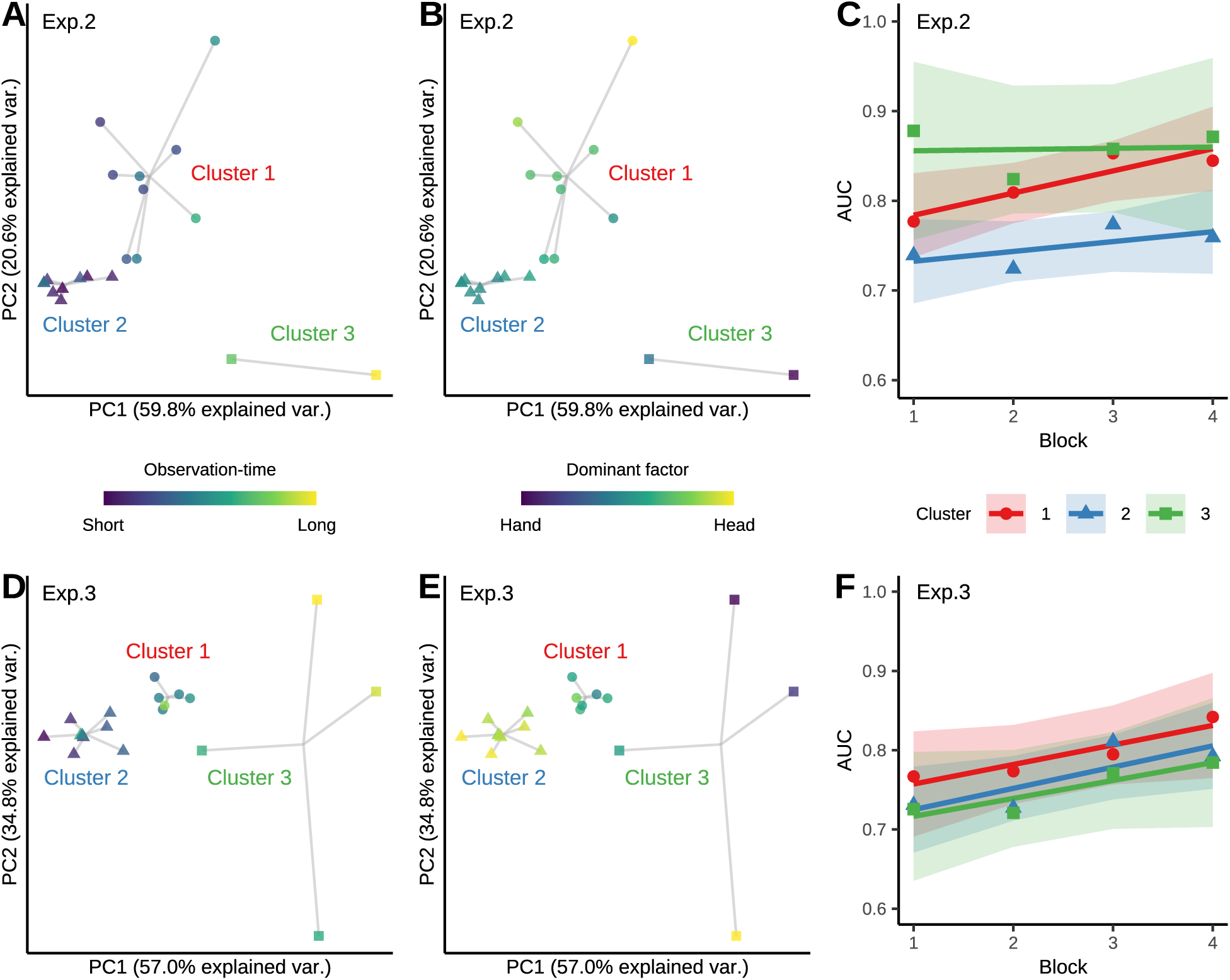
Individual differences among the exploratory strategies used in Experiments 2 and 3. (A–B) Results of PCA applied to participants’ exploratory behaviors in Experiment 2. Each point represents one participant, and the marker shapes denote cluster membership. Points are color-coded according to their (A) overall observation times and (B) relative reliance on head movements versus hand movements. (C) Discrimination performance (AUC) changes observed across different experimental blocks for each PCA-defined cluster in Experiment 2. The experiment was divided into four successive blocks from beginning to end. Shaded regions indicate 95% confidence intervals around the regression lines. (D–F) PCA results for the exploratory behaviors observed in Experiment 3. The format is identical to that employed in (A–C).

Similarly, in Experiment 3, PCA followed by GMM clustering revealed a two-dimensional structure underlying the individual exploratory behavior differences, indicating that comparable factors accounted for the variability observed across different participants (Figures 4D and 4E). In contrast with those in Experiment 2, however, the differences among the learning trajectories produced across various clusters were less pronounced in Experiment 3 (*F* (2, 16) = 0.77, *p* = 0.478, *η*^2^ = 0.088; Figure 4F), whereas the overall discrimination performance improved the across blocks in all groups (*F* (1, 54) = 7.19, *p* = 0.010, *η*^2^ = 0.118). This pattern suggests that although the participants relied on distinct exploratory strategies to acquire diagnostic cues, they were nonetheless able to enhance their performance over time. Together, these results highlight the flexibility of active information sampling, showing that observers can converge on effective material discrimination performance even when they adopt different exploration strategies.

## Discussion

A large body of work has established the foundations of material perception, describing how mid-level visual processes recover surface properties from the interplay of shading, specular reflections, and three-dimensional shapes (1–5). These frameworks highlight the fact that many material cues are inherently dynamic, varying systematically with changes in the viewpoint, object pose, and illumination level. Many of these dynamic cues, however, have been characterized using stimuli in which motion is generated by the experimenter rather than the observer. Such viewpoint-dependent transformations—including changes in specular flow, highlight motion, and shading changes—provide powerful information for discriminating among material categories.

However, most of this work has relied on passive or experimenter-controlled optical structure changes, raising the question of how observers actively select actions to acquire diagnostic cues.

### Active Viewing Fundamentally Reshapes Material Recognition

Our three experiments demonstrate that active viewing fundamentally shapes the human material recognition process. Allowing observers to freely move their heads and hands increased the flexibility with which they sampled diagnostic visual information, and their discrimination performance improved when these exploratory actions yielded informative cues (Figures 1G and 1H). By dissociating the contributions of viewpoint changes and object manipulations, we found that head motion enhanced the discrimination results when comparative cues were available (Figure 2G), whereas hand-based object manipulations played a dominant role when viewpoint cues were limited (Figure 3C). In both cases, greater use of the modality that provided useful information was associated with higher discrimination accuracy (Figures 2I and 3E). Clustering analyses further revealed that participants who engaged in more efficient sampling strategies achieved greater performance gains over time (Figure 4C) and demonstrated notable flexibility in terms of enhancing their discrimination accuracy even when they relied on distinct exploratory strategies (Figure 4E). Together, these findings reveal that material recognition is supported by context-dependent exploratory strategies—an aspect of visual perception that is largely undetectable under conventional passive-viewing paradigms.

Several recent studies have emphasized the claim that active explorations can enhance material perception capabilities. For example, Lin et al. (2025) reported that allowing observers to freely manipulate objects improved their perceptual judgments, with exploratory movements being particularly extensive—and presumably more informative—during gloss judgments relative to lightness judgments (25). However, their findings did not specify when active exploration becomes beneficial, which exploratory modalities drive this improvement, or how these contributions depend on the viewing context. By systematically separating head- and hand-based cues and comparing conditions in which viewpoint or manipulation information was more or less useful, our experiments demonstrated that the effectiveness of active exploration is highly context-dependent.

Crucially, the number of exploratory movements increased specifically under conditions of heightened sensory ambiguity, indicating that observers selected actions to reduce their perceptual uncertainty. This interpretation suggests that the visual system flexibly adapts its sampling strategy to obtain the most diagnostic information available—an idea consistent with theories concerning adaptive, goal-directed perception (26). Notably, when one modality was disabled, the participants often produced “empty” head or hand movements that did not influence the stimulus (Figures 3F and 3G), revealing a strategic intent to seek missing cues even when such movements had no visual consequences. This pattern is in line with the perception–action loop framework, which holds that exploratory behaviors constitute an integral part of perceptual inference rather than ancillary motor outputs.

### Context-Dependent Contributions of Head- and Hand-Based Explorations

How do observers determine which exploratory action to use when identifying a material? Across our experiments, head motions and hand motions contributed to the discrimination process in different viewing contexts, suggesting that each action generates distinct diagnostic cues. The viewpoint changes produced by head movements yield specular-flow and motion-parallax information, which becomes particularly informative when multiple objects can be compared simultaneously (16–18). In contrast, object manipulation alters the pose of the target object and reveals different highlighting, refractive, and reflective configurations (19–24), making hand-based exploration more effective when viewpoint cues are impoverished. Together, these patterns suggest that observers flexibly evaluate the informativeness of the available dynamic cues and select the exploratory action that best reduces the degree of perceptual uncertainty in a given context.

### Illumination Structure, Individual Differences, and Learning in Active Material Perception Cases

Even when the material category remained identical, the observers’ responses varied substantially across differenct illumination environments, which was consistent with prior work showing strong lighting-dependent material appearance modulation (14, 15). Our experiments included not only natural illumination maps captured from real-world scenes but also an artificial checker-pattern light field (Figures 2E and S1G). Under natural lighting, observers could exploit priors such as the light-from-above assumption (41, 42). Under artificial illumination, their performance was reduced relative to that attained under natural-light conditions; however, the observers were nevertheless able to perform well above random chance, although a purely static cue would have made the task nearly impossible (23). These results indicate that, regardless of the structure of the illumination environment, the dynamic information generated by viewpoint changes and object manipulations provides the principal cues for resolving material identities.

Quantifying the participants’ movements allowed us to reveal individual differences in how efficiently the observers extracted diagnostic cues during active explorations (Figure 4). Under traditional passive-viewing paradigms, all observers receive identical retinal inputs, limiting the extent to which individual material perception variability can be expressed. In contrast, active sensing enables substantial performance differences because observers differ not only in their low-level sensory abilities but also in the higher-level strategies that they deploy when selecting and integrating cues (43–45). Because our participants were young adults, the observed differences likely reflect strategic sampling variations rather than age-related sensory declines. However, the origins of these strategy differences remain unclear.

Moreover, because these exploratory strategies were also associated with the magnitude of learning across different trials, individual perceptual learning efficiency differences may also have contributed to the observed performance variability. Alongside theoretical proposals that stipulate that material perception emerges from data-driven learning over development and that humans can often generalize to new materials from very small numbers of exemplars, this pattern suggests that both long-term experience and relatively rapid learning processes may shape how efficiently observers discover and exploit diagnostic cues (2).

### Limitations and Directions for Future Research

One limitation of this research pertains to the range of materials tested. The present study focused exclusively on metal and glass. At first glance, this restriction may appear to limit the generalizability of our conclusions. However, distinguishing metal from glass is among the most challenging problems in the material perception domain because the appearances of both materials depend strongly on the associated environmental illumination conditions and arises from the complex interplay between specular reflection and refraction (8, 14, 23, 24, 27, 28). For precisely this reason, examining how active exploration contributes to discriminating between these two optically demanding categories offers a stringent test of the mechanisms underlying material recognition from a broader perspective. Nevertheless, the extent to which the context-dependent sampling strategies observed here can be generalized to other material classes—such as wood, fabric, plastic, or liquid—remains unclear. Future work should expand the stimulus space to include materials with diverse reflectance, refractive, and subsurface scattering properties to identify the boundary conditions and general domain principles of active material perception. Large-scale analyses of material judgments indicate that diverse materials can be described by a compact set of shared perceptual dimensions (46), suggesting that some of the cues leveraged in metal–glass discrimination tasks may also support judgments of other material classes. Nevertheless, direct tests will be required to determine the breadth of this generalization.

Moreover, the present study focused on rigid objects, but many real-world materials do not maintain a fixed shapes, and how observers actively interact with such materials remains largely unknown. Even nominally solid objects can deform under active manipulations, revealing elasticity or stiffness cues (47–49). Other materials exhibit viscosity or fluid-like behaviors and lack stable geometries (50–53). In addition, observers often form predictions about the likely material properties of an object before interacting with it (54, 55), and some materials may even fracture or break upon contact (56). By demonstrating how active exploration supports the discrimination of rigid materials, the present framework provides a foundation for extending active-sensing approaches to these broader classes of deformable, fluid, and fragile materials.

A second limitation concerns the use of virtual reality. Although VR allowed us to experimentally dissociate head and hand movements—two sources of information that are tightly coupled in natural behaviors—this very dissociation is artificial in real-world settings. Moreover, the ability to manipulate these modalities independently is a key strength of VR, enabling us to demonstrate that observers flexibly reweight their sampling strategies depending on which modality provides more diagnostic information. It remains an open question whether the same context-dependent strategy switching emerges when head and hand movements are intrinsically coupled in the physical world. A related question concerns the ecological validity of providing simultaneous reference objects, as done in Experiment 2. Although real environments contain many materials, situations requiring observers to directly compare multiple surfaces with high reflectance or transparency levels may be relatively rare. In contrast, scenarios that more closely resembles Experiment 3, where a single target object must be judged among a heterogeneous set of background materials, may capture the material identification problems that are typically faced in everyday vision settings more effectively.

Furthermore, real-world material perception is an inherently multisensory process (57, 58). The haptic feedback obtained when interacting with an object (e.g., surface compliance, roughness, or temperature (59–61)) and physical cues such as weight or thermal conductivity can substantially influence material judgments. Although our VR setup included minimal vibrotactile feedback when the virtual hand contacted the object to enhance the degree of immersion, it did not provide any diagnostic haptic information that was relevant to the material properties. Thus, the present study effectively isolated the visual component and demonstrated that observers deploy context-dependent strategies based primarily on visual explorations. This framework provides a foundation for future researchers to investigate how active visual sampling can be integrated with richer haptic and proprioceptive signals during naturalistic material recognition processes.

A third limitation concerns the acquisition of active sampling strategies. Our results showed that individual differences in how the participants explored the stimulus—such as pulling the object, approaching it, or viewing it from below or above—were systematically related to both the material identification accuracy and learning rate achieved across different trials (Figure 4). This suggests that the visual system benefits from efficient exploration strategies, yet how such strategies are acquired remains unknown. Do observers possess innate preferences for certain visual or motion cues, or are these strategies learned through experience? Notably, even within the few hundred trials of our experiment, some participants appeared to implicitly (or explicitly) discover more efficient motion patterns and subsequently improved their performance. This raises the possibility that strategy discovery and refinement occur on relatively short timescales. One promising extension of the present study is to analyze movement trajectories and viewpoint sequences in greater detail—linking motion features and video-based appearance cues to trial-by-trial decision outcomes.

Computational modeling may also help identify which cues an ideal observer would exploit and whether human observers converge on similar strategies (6–13), thereby clarifying which aspects of active material perception reflect learned behaviors versus the inherent constraints imposed on the visual system.

In summary, our findings demonstrate that human observers flexibly adapt their active sampling strategies to the given viewing context constraints, enabling both higher material identification accuracy and faster performance gains to be achieved. These results advance a theoretical framework in which material perception is understood not as a passive static image decoding procedure but rather as an adaptive process shaped by task demands and exploratory behaviors, revealing aspects of material perception that remain inaccessible under the traditional passive-viewing paradigms.

## Materials and Methods

### Participants

Twenty students at the Toyohashi University of Technology participated in each of Experiments 1, 2 and 3. The sample size was determined on the basis of previous virtual-reality material perception studies, which typically employed between 12 and 20 participants (62, 63). All the participants had normal or corrected-to-normal vision and were naive to the purpose of the experiments. Four individuals participated in both Experiments 1 and 2, whereas there was no overlap between Experiments 2 and 3. All participants were provided with a full explanation of the experimental procedures and provided written informed consent prior to participating. This study was approved by the Committee for Human Research of the Toyohashi University of Technology.

### Apparatus

The participants were seated in a chair and wore a Varjo Aero head-mounted display (resolution: 2880 × 2720 per eye; field of view: 115°; refresh rate: 90 Hz). They additionally held a pair of VR controllers (HTC Vive controller), one in each hand. Positional tracking was achieved using SteamVR Base Station 2.0. The virtual environments were implemented in Unity (version 2022.3.20f1 for Experiment 1 and 2022.3.22f1 for Experiments 2 and 3) with the Varjo XR Plugin (version 3.7.0) and SteamVR Plugin (version 2.8.0). To perform physically accurate rendering, we used Unity’s High-Definition Rendering Pipeline (HDRP). To minimize the degree of simulator sickness, it was essential to maintain a smooth rendering process with minimal latency and flicker. Therefore, we adopted a BSDF-based material model. Because the primary visual cues of interest in this study concerned surface reflection and transmission, subsurface scattering was not needed. This modeling choice allowed us to achieve stable, high-fidelity rendering while maintaining high real-time performance.

### Stimuli

A randomly bumped three-dimensional object (8 cm × 8 cm × 8 cm) was presented at a viewing distance of 25 cm in Experiment 1. In Experiments 2 and 3, a geometrically identical but smaller object (6 cm × 6 cm × 6 cm) was presented at a viewing distance of 50 cm. At these initial spatial arrangements, the objects subtended visual angles of 18.19° (Experiment 1) and 6.87° (Experiments 2 and 3), respectively. The materials were implemented using the HDRP Lit shader in Unity, with the following parameters fixed: smoothness = 1.0, index of refraction = 1.424, and thickness = 0.8. The metallic parameter determined the degree to which the surface appeared metallic. By varying this parameter from 0.0 to 1.0, we generated a continuous transition from a glass-like appearance to a metal-like appearance. This parameter served as the metallicity factor. In Experiments 1 and 2, seven levels were used (0.0, 0.17, 0.33, 0.50, 0.67, 0.83, and 1.0), whereas Experiment 3 employed five levels (0.0, 0.25, 0.5, 0.75, and 1.0). Thus, the stimuli spanned a continuum from transparent glass-like objects to mirror-like metallic objects. Subsurface scattering was disabled to avoid unintended translucent appearances at intermediate levels (see also the “Apparatus” section). To validate the perceptual plausibility of these materials, we conducted a matching experiment in which the participants compared our stimuli to physically based renderings with systematically varied refractive indices (23). The results (*n* = 7) confirmed that stimuli with higher metallicity levels were consistently matched to materials with higher refractive indices (Figure S2C).

Two object shapes were used on the basis of prior work (23): a relatively smooth object (“smooth”) and a more irregular object (“bumpy”), which was created by randomly modulated surface perturbations (see Figure S1A).

Illumination was provided using 4*π*-steradian high-dynamic-range-imaging (HDRI) environment maps obtained from Poly Haven (https://polyhaven.com/, CC0 license). In Experiment 1, two outdoor (Sky and Rainforest) and two indoor (Studio and Empty Room) lighting environments were selected from this database (Figure S1B). In addition, two artificial environments generated in Blender were used, including one structured environment (Checkerboard) and one unstructured environment (Wave Noise). Experiments 2 and 3 employed two natural HDRI environments (Sky and Studio) and two artificial environments (Check and Gray) (the illumination conditions are shown in Figure S1C; example stimuli are presented in Figure S1D).

## Procedure

### Experiment 1

Each participant sat on a chair while wearing the HMD and holding a controller in each hand. A target object was presented 25 cm in front of the participant. Two reference objects—one fully glass object (metallicity = 0.0) and one fully metal object (metallicity = 1.0)—were placed 75 cm to the left and right of the target (Figure 1C), respectively. This arrangement prevented the target and the reference objects from being visible simultaneously. The participants used the trigger on the right-hand controller to virtually grasp the target object and freely translate and rotate it. Hand movements were rendered in real time using a glove-shaped virtual avatar linked to the controller. To provide haptic confirmation during manipulations, the controller delivered a brief vibration whenever the virtual hand made contact with the object. The task was to judge whether the material of the central target was closer to the glass or metal reference, using a slider bar controlled by the left-hand controller. Participants were required to observe the stimulus for at least 6 s before they responded. Responses were disabled during this initial period to prevent immediate judgments based solely on a single static view. This duration was determined on the basis of pilot testing to allow sufficient time for exploratory interactions. Each participant completed 168 trials, including metallicity (7), shape (2), illumination (6), and reference position (2) trials. The participants were allowed to take a break following every 42 trials. Five practice trials were conducted before the main experiment.

### Experiment 2

The target stimulus was presented 50 cm in front of the participant and was flanked by two reference stimuli positioned 17 cm to the left and right (Figure 2A). The participants compared the target with the two reference stimuli and responded by indicating which reference it most closely resembled using a button located on the left-hand controller. The degree of observational freedom was manipulated by orthogonally combining the presence or absence of head motions (HMD tracking enabled or disabled, respectively) and hand motions (controller tracking enabled or disabled, respectively), yielding four experimental conditions (Figure 2B). Under the hand-motion-available conditions, the participants could grasp and freely manipulate the target stimulus with their right hands. The reference stimuli were either fully glass-like (metallicity = 0.0) or fully metal-like (metallicity = 1.0), and the left–right assignment of these reference stimuli was counterbalanced and treated as the reference position factor. Each participant completed 224 trials in one day, comprising all combinations of the following factors: the observational condition (4), metallicity level (7), shape (2), reference position (2), and illumination level (2). The trials were blocked by the observational condition, with each block consisting of 56 trials, resulting in four blocks per participant. The order of the blocks was counterbalanced across the participants, and the order of the trials contained within each block was randomized. The experiment was conducted across two separate days. Among the four illumination conditions (Figure S1C), the two natural illumination conditions were tested on one day, and the two artificial illumination conditions were tested on the other day. Thus, each participant completed a total of 448 trials. The order of these sessions was counterbalanced across the participants. The participants completed a short practice session before the main experiment, consisting of eight trials formed by the combination of the observational condition (4) and illumination level (2) in each session.

### Experiment 3

The procedure used for this experiment was almost identical to that of Experiment 2, except that the task changed from judging which reference object the target more closely resembled to judging whether the target appeared more like metal or glass (Figure 3A). The experimental session consisted of 160 trials in total; this number was derived from the full factorial combination of the observational condition (4), metallicity (5), shape (2), and illumination level (4). The trials were blocked by the observational conditions, with each block containing 40 trials, resulting in four blocks in total. The order of the blocks was counterbalanced across the participants, and the trials within each block were presented in a randomized order. After completing the practice trials, the participants proceeded to the main experiment. The practice trials consisted of 16 trials, corresponding to all combinations of observational conditions (4) and illumination levels (4).

In Experiment 3, we additionally recorded head and hand movements even when head tracking or hand tracking was disabled. Under these “tracking-off” conditions, movements made by the HMD or controller did not affect the visual scene, meaning that the stimulus did not respond to the participants’ actions. Nevertheless, the participants often moved their heads or hands, and these “empty” exploratory movements were fully logged for analysis purposes (see also Figures 3F and 3G).

## Data Analysis

All analyses were conducted in R (version 4.5.1). In Experiment 1, two participants were excluded because they responded in a binary manner for a substantial portion of the session, indicating that they did not fully understand the continuous slider response system. The final analysis therefore included 18 participants. No participants were excluded from Experiment 2 (*n* = 20). In Experiment 3, one participant was removed because of their inability to obtain proper optical focus while wearing the HMD, resulting in a final sample consisting of 19 participants.

### Response Data

**Experiment 1.** For each trial, the participants rated how similar the target stimulus appeared relative to each of the two reference stimuli. Accuracy was computed by normalizing the absolute difference between each participant’s rating and the true material value (metallicity) of the stimulus by the maximum possible difference. The accuracy scores were then averaged separately for each participant (Figure 1H).

For each participant and condition, the corresponding metallicity rating was analyzed using a linear mixed-effects model implemented in the lme4 package (version 1.1.37). The model included the metallicity rating *R* as the dependent variable, with metallicity *M* and illumination *E* as fixed effects and as an interaction term, and the participant ID and shape *G* as random effects. The model was specified as: *R* ∼ *M* ∗ *E* + (1 + *M* |*ID*) + (1|*G*). Effect sizes are reported as likelihood-based estimates of partial eta squared (*η*^2^) values for the linear mixed-effects model (Figure 1F), which were computed using the eta_squared function from the effectsize package.

**Experiments 2 and 3.** To quantify the discriminability of materials under each observation condition (Figures 2G and 3C), we computed the area under the receiver operating characteristic curve (AUC) using the pROC package (version 1.19.0.1). We then performed a repeated-measures two-factor analysis of variance (ANOVA), treating the presence or absence of head and hand movements as within-subject factors. The normality of the AUC values within each condition was assessed using the Shapiro–Wilk test. Because the assumption of normality was violated in Experiment 2, we applied an aligned rank transform (ART) using the ARTool package (version 0.11.2) and conducted a two-way ANOVA on the transformed data. Notably, effect sizes such as partial eta squared were not reported because the ART operates on rank-transformed data and does not preserve variance information in the original measurement scale. As a result, variance-based effect size measures are not directly interpretable for ART-based ANOVA.

### Behavioral Data

**Translational Movement.** For each condition, we computed the across-participant mean movement distances of the head and right hand. For each trial, the total movement distance *D* was calculated using the following equation:

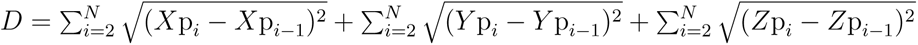 where *N* denotes the number of frames within a trial, and *X*p*_i_, Y* p*_i_, Z*p*_i_* represent the XYZ coordinates of either the HMD (head) or the experimental stimulus (hand) at frame *i*. The amount of translation was then averaged within each observation condition and used in subsequent analyses.

**Rotational Movement.** We computed the mean rotational displacements of the HMD (head) and of the stimulus (hand). For a given trial, the rotational displacement *R* was quantified according to the following equation:

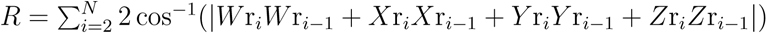, where *N* denotes the number of frames within a trial and *W* r*_i_, X*r*_i_, Y* r*_i_, Z*r*_i_* represent the quaternion components of the HMD (or of the stimulus) at frame *i*. The resulting rotational displacement values were averaged within each observation condition for further analysis.

**Polynomial Modeling of Exploratory Behavior as a Function of Material Ambiguity.** To determine whether the participants modulated their exploratory behaviors as a function of material ambiguity, we predicted translational (or rotational) movement using a polynomial regression model as a function of the metallicity of the stimulus. On the basis of the hypothesis that participants would engage in more active information sampling for ambiguous stimuli (e.g., metallicity = 0.50) than for unambiguous stimuli (e.g., metallicity = 0.0 or 1.0), the translational distance *D* was modeled as a function of the linear and quadratic metallicity terms *M* : *D* ∼ *M* + *M* ^2^. Similarly, rotational movement *R* was modeled using the same polynomial structure: *R* ∼ *M* + *M* ^2^.

### Individual Exploratory Strategy Differences

**Classification of Exploratory Behaviors.** The participants’ exploratory behaviors were categorized into four types, and the time spent in each category was quantified (Figures S2A and S2B). “Pulling the stimulus” was defined as instances in which the stimulus–head distance was 40 cm or less and the stimulus had moved farther from its initial position than the participant’s head, indicating that the object was actively drawn toward the observer. “Approaching the stimulus” was identified when the stimulus–head distance was likewise within 40 cm, but the participant’s head had moved farther from its initial position than the stimulus, which was consistent with self-motion toward the object rather than object manipulation. “Viewing from below” was assigned when the stimulus was positioned more than 5 cm above eye level, indicating observation sfrom underneath.

Conversely, “viewing from above” was defined as occuring when the participant’s head was more than 5 cm higher than the stimulus, reflecting observations from an elevated viewpoint.

**Dimensionality Reduction and Clustering of Exploratory Strategies.** To identify the principal dimensions underlying individual differences in exploratory behavior differences, we performed PCA on the time spent in each of the four exploratory categories defined above separately for Experiments 2 and 3. The first two principal components were retained for further analysis. Gaussian mixture model (GMM) clustering was then applied to the PCA scores to classify the participants into distinct groups on the basis of their exploratory strategies. The optimal number of clusters was determined to be three for both Experiments 2 and 3.

The exploratory strategies were visualized on the basis of the scores of the first and second principal components (Figures 4A and 4B for Experiment 2; Figures 4D and 4E for Experiment 3). The data points were color-coded according to (1) the overall observation time and (2) the relative reliance on head-based versus hand-based exploration, which were quantified from both sets of raw observation data (Figures S2A and S2B).

To examine the learning effects induced across clusters, the discrimination accuracy (AUC) was computed for each participant in each of four successive trial blocks obtained by dividing the full experiment into quartiles. Changes in the AUC across different blocks were then analyzed using a linear mixed-effects model. The AUC was entered as the dependent variable, with blocks and clusters as fixed effects and participant as a random intercept. The model was defined as follows: *A* ∼ *B* ∗ *C* + (1|*ID*), where *A* denotes the AUC, *B* denotes the block, and *C* denotes the cluster.

## Acknowledgments

This work was supported by JSPS KAKENHI (Grant Numbers JP25K21323 to H.T., JP25H01141 to S.N., and JP23KK0183 to T.M.) and the Toukai Foundation for Technology.

Conflicts of Interest

The authors declare that they have no competing interests.

## Data and Code Availability

The datasets and analysis code that support the main findings of this study are available at the Open Science Framework repository (https://osf.io/xzunv/). Additional raw motion-tracking data that are still part of ongoing analyses by our group will be made available upon reasonable request to the corresponding author.

## Declaration of Generative AI and AI-Assisted Technologies during the Writing Process

During the preparation of this work, the authors used ChatGPT 5.1 to improve the language, and the manuscript has been proofread by native English speakers through an English editing service. Following this use of the tool and service, the authors reviewed and edited the content as needed and take full responsibility for the content of the publication.

## Supplemental Information: Supplemental Figures

**Figure S1.**
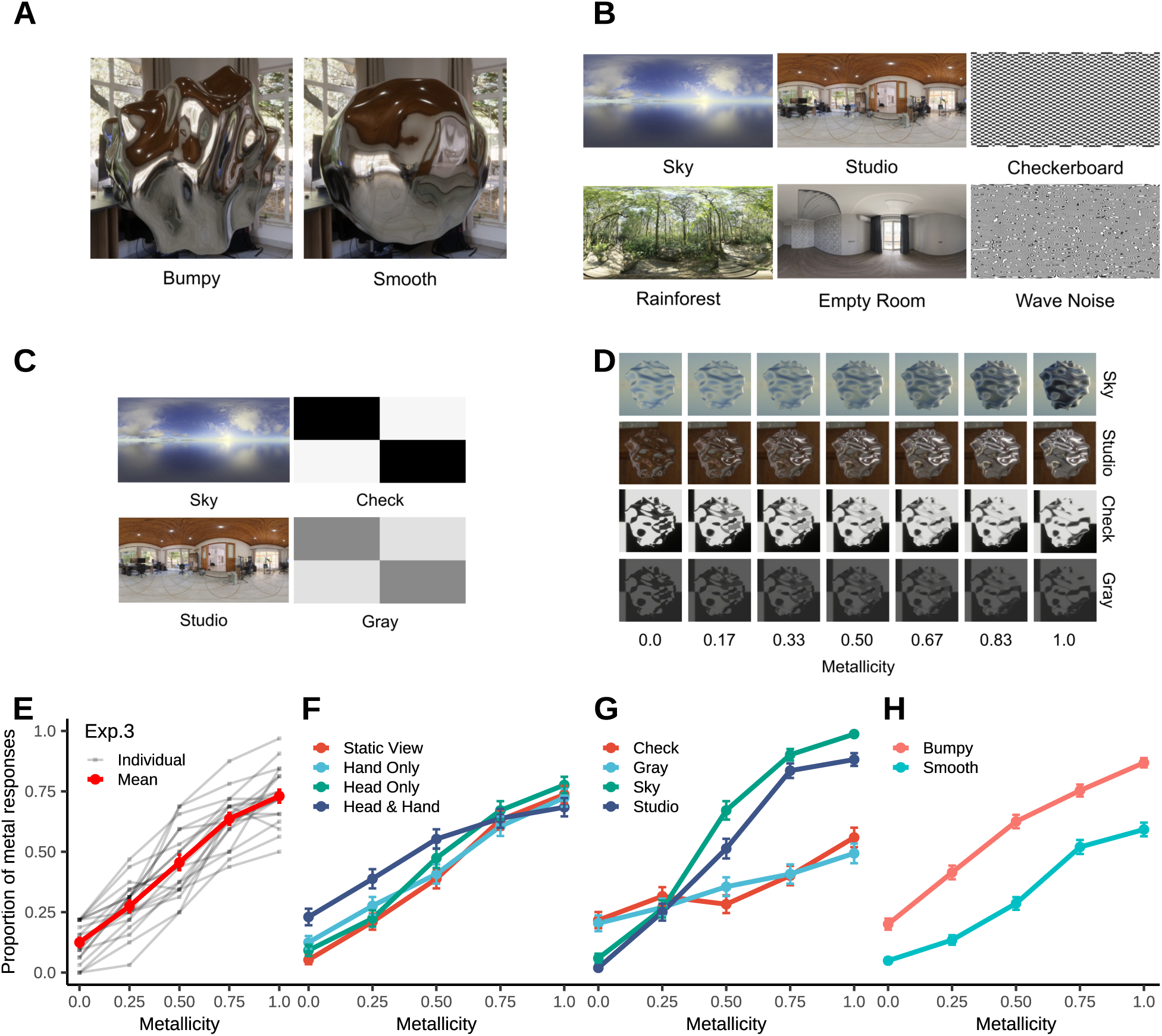
(A) Examples of stimulus shapes: a highly bumpy object (left) and a relatively smooth object (right). (B) Examples of the six illumination environments used in Experiment 1. Columns from left to right: outdoor, indoor, and artificial environments. (C) Examples of the four illumination environments used in Experiments 2 and 3. From left to right, the columns denote natural and artificial environments. (D) Examples of the stimuli used in Experiments 2 and 3, illustrating the material appearance manipulations made along the metallicity parameter from 0.0 (left) to 1.0 (right). (E–H) Metal response data obtained in Experiment 3: overall and individual results (E), plotted separately by experimental condition (F), illumination environment (G), and object shape (H).

**Figure S2.**
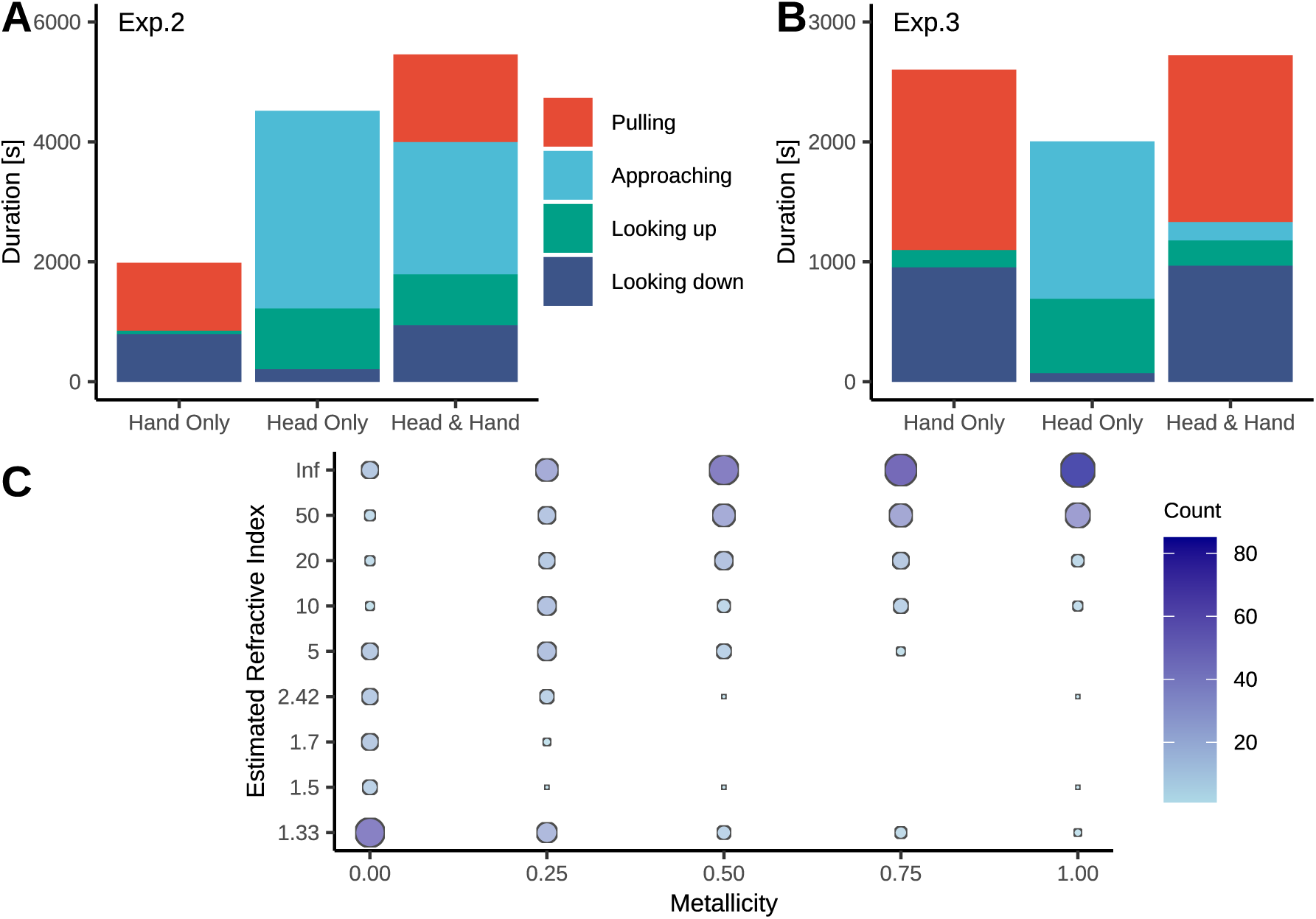
(A) Total exploration time required for all participants in Experiment 2. (B) Total exploration time required for all participants in Experiment 3. (C) Results of a matching experiment showing the correspondence between the metallicity parameter and the refractive index obtained from the physically based rendering procedure (n = 7).

## References

1. R. W. Fleming, Visual perception of materials and their properties. Vision Research 94, 62–75 (2014).

2. R. W. Fleming, Material Perception. Annual Review of Vision Science 3, 365—388 (2017).

3. B. L. Anderson, Visual perception of materials and surfaces. Current Biology 21, R978–R983 (2011).

4. B. L. Anderson, Mid-level vision. Current Biology 30, R105–R109 (2020).

5. B. L. Anderson, P. J. Marlow, Perceiving the shape and material properties of 3D surfaces. Trends in Cognitive Sciences 27, 98–110 (2023).

6. K. E. Prokott, H. Tamura, R. W. Fleming, Gloss perception: Searching for a deep neural network that behaves like humans. Journal of Vision 21, 1–20 (2021).

7. E. Prokott, R. W. Fleming, Identifying specular highlights: Insights from deep learning. Journal of Vision 22, 1–19 (2022).

8. H. Tamura, K. E. Prokott, R. W. Fleming, Distinguishing mirror from glass: A “big data” approach to material perception. Journal of Vision 22, 1–22 (2022).

9. K. R. Storrs, B. L. Anderson, R. W. Fleming, Unsupervised learning predicts human perception and misperception of gloss. Nature Human Behaviour 5, 1402–1417 (2021).

10. C. Liao, M. Sawayama, B. Xiao, Unsupervised learning reveals interpretable latent representations for translucency perception. PLOS Computational Biology 19, e1010878 (2023).

11. J. R. Cheeseman, J. A. Ferwerda, T. Morimoto, R. W. Fleming, Gloss discrimination: Toward an image-based perceptual model. Journal of Vision 25, 1–19 (2025).

12. J. J. R. van Assen, S. Nishida, R. W. Fleming, Visual perception of liquids: Insights from deep neural networks. PLOS Computational Biology 16, e1008018 (2020).

13. T. Morimoto, et al., Human gloss perception reproduced by tiny neural networks. bioRxiv 2025.05.09.653112 (2025). 10.1101/2025.05.09.653112.

14. J. Kim, P. J. Marlow, Turning the World Upside Down to Understand Perceived Transparency. i-Perception 7, 1—5 (2016).

15. B. Xiao, et al., Looking against the light: How perception of translucency depends on lighting direction. Journal of Vision 14, 1–22 (2014).

16. Y. Sakano, H. Ando, Effects of head motion and stereo viewing on perceived glossiness. Journal of Vision 10, 1–14 (2010).

17. Y. Tani, et al., Enhancement of Glossiness Perception by Retinal-Image Motion: Additional Effect of Head-Yoked Motion Parallax. PLoS ONE 8, e54549 (2013).

18. P. J. Marlow, B. L. Anderson, Motion and texture shape cues modulate perceived material properties. Journal of Vision 16, 1—14 (2016).

19. K. Doerschner, et al., Visual Motion and the Perception of Surface Material. Current Biology 21, 2010—2016 (2011).

20. G. Wendt, F. Faul, V. Ekroll, R. Mausfeld, Disparity, motion, and color information improve gloss constancy performance. Journal of Vision 10, 1—17 (2010).

21. J. Kim, P. Marlow, B. L. Anderson, The perception of gloss depends on highlight congruence with surface shading. Journal of Vision 11, 1–19 (2011).

22. K. Doerschner, O. Yilmaz, G. Kucukoglu, R. W. Fleming, Effects of surface reflectance and 3D shape on perceived rotation axis. Journal of Vision 13, 1–23 (2013).

23. H. Tamura, H. Higashi, S. Nakauchi, Dynamic Visual Cues for Differentiating Mirror and Glass. Scientific Reports 8, 8403 (2018).

24. M. Ohara, J. Kim, K. Koida, The Effect of Material Properties on the Perceived Shape of Three-Dimensional Objects. i-Perception 11, 1–14 (2020).

25. L. P. Y. Lin, K. Drewing, K. Doerschner, Exploring for gloss: Active exploration in visual material perception. Journal of Vision 25, 1–17 (2025).

26. B. Xiao, C. Liao, Material perception connects vision, cognition and action. Nature Reviews Psychology 4, 687–701 (2025).

27. H. Tamura, S. Nakauchi, The Rotating Glass Illusion: Material Appearance Is Bound to Perceived Shape and Motion. i-Perception 9, 1—5 (2018).

28. M. Ohara, J. Kim, K. Koida, The Role of Specular Reflections and Illumination in the Perception of Thickness in Solid Transparent Objects. Frontiers in Psychology 13, 766056 (2022).

29. R. W. Fleming, A. Torralba, E. H. Adelson, Specular reflections and the perception of shape. Journal of Vision 4, 10 (2004).

30. A. A. Muryy, R. W. Fleming, A. E. Welchman, “Proto-rivalry”: how the binocular brain identifies gloss. Proceedings of the Royal Society B: Biological Sciences 283, 20160383 (2016).

31. J. T. Todd, J. F. Norman, The visual perception of metal. Journal of Vision 18, 1–17 (2018).

32. J. S. Harvey, H. E. Smithson, Low level visual features support robust material perception in the judgement of metallicity. Scientific Reports 11, 16396 (2021).

33. A. C. Schmid, P. Barla, K. Doerschner, Material category of visual objects computed from specular image structure. Nature Human Behaviour 7, 1152–1169 (2023).

34. R. W. Fleming, F. Jakel, L. T. Maloney, Visual Perception of Thick Transparent Materials. Psychological Science 22, 812—820 (2011).

35. N. Schlüter, F. Faul, Are optical distortions used as a cue for material properties of thick transparent objects? Journal of Vision 14, 1—14 (2014).

36. N. Schlüter, F. Faul, Matching the Material of Transparent Objects: The Role of Background Distortions. i-Perception 7, 1—24 (2016).

37. N. Schlüter, F. Faul, Visual shape perception in the case of transparent objects. Journal of Vision 19, 1–36 (2019).

38. J. T. Todd, J. F. Norman, Reflections on glass. Journal of Vision 19, 1–21 (2019).

39. J. T. Todd, J. F. Norman, Contours produced by internal specular interreflections provide visual information for the perception of glass materials. Journal of Vision 20, 1–18 (2020).

40. D. Gigilashvili, J.-B. Thomas, J. Y. Hardeberg, M. Pedersen, Translucency perception: A review. Journal of Vision 21, 1–41 (2021).

41. W. J. Adams, E. W. Graf, M. O. Ernst, Experience can change the ’light-from-above’ prior. Nature Neuroscience 7, 1057–1058 (2004).

42. W. J. Adams, et al., The Southampton-York Natural Scenes (SYNS) dataset: Statistics of surface attitude. Scientific Reports 6, 35805 (2016).

43. J. F. Norman, et al., Aging and the Haptic Perception of Material Properties. Perception 45, 1387–1398 (2016).

44. M. O. Ernst, M. S. Banks, Humans integrate visual and haptic information in a statistically optimal fashion. Nature 415, 429–433 (2002).

45. L. T. Maloney, H. Zhang, Decision-theoretic models of visual perception and action. Vision Research 50, 2362–2374 (2010).

46. F. Schmidt, M. N. Hebart, A. C. Schmid, R. W. Fleming, Core dimensions of human material perception. Proceedings of the National Academy of Sciences 122, e2417202122 (2025).

47. V. C. Paulun, F. Schmidt, J. J. R. van Assen, R. W. Fleming, Shape, motion, and optical cues to stiffness of elastic objects. Journal of Vision 17, 1—22 (2017).

48. F. Schmidt, V. C. Paulun, J. J. R. van Assen, R. W. Fleming, Inferring the stiffness of unfamiliar objects from optical, shape and motion cues. Journal of Vision 17, 1—17 (2017).

49. V. C. Paulun, R. W. Fleming, Visually inferring elasticity from the motion trajectory of bouncing cubes. Journal of Vision 20, 1–14 (2020).

50. T. Kawabe, K. Maruya, S. Nishida, Perceptual transparency from image deformation. Proceedings of the National Academy of Sciences of the United States of America 112, E4620—E4627 (2015).

51. T. Kawabe, K. Maruya, R. W. Fleming, S. Nishida, Seeing liquids from visual motion. Vision Research 109, 125—138 (2015).

52. J. J. R. van Assen, R. W. Fleming, Influence of optical material properties on the perception of liquids. Journal of Vision 16, 1—20 (2016).

53. J. J. R. van Assen, P. Barla, R. W. Fleming, Visual Features in the Perception of Liquids. Current Biology 28, 452–458 (2018).

54. L. M. Alley, A. C. Schmid, K. Doerschner, Expectations affect the perception of material properties. Journal of Vision 20, 1–20 (2020).

55. D. Kaiser, R. Stecher, K. Doerschner, EEG Decoding Reveals Neural Predictions for Naturalistic Material Behaviors. The Journal of Neuroscience 43, 5406–5413 (2023).

56. A. C. Schmid, K. Doerschner, Shatter and splatter: The contribution of mechanical and optical properties to the perception of soft and hard breaking materials. Journal of Vision 18, 1–32 (2018).

57. W. Fujisaki, N. Goda, I. Motoyoshi, H. Komatsu, S. Nishida, Audiovisual integration in the human perception of materials. Journal of Vision 14, 1–20 (2014).

58. C. Spence, Shitsukan — the Multisensory Perception of Quality. Multisensory Research 33, 737–775 (2020).

59. S. Okamoto, H. Nagano, Y. Yamada, Psychophysical Dimensions of Tactile Perception of Textures. IEEE Transactions on Haptics 6, 81–93 (2013).

60. D. N. Dövenciglu, F. S. Üstün, K. Doerschner, K. Drewing, Hand explorations are determined by the characteristics of the perceptual space of real-world materials from silk to sand. Scientific Reports 12, 14785 (2022).

61. L. A. Jones, H.-N. Ho, Tactile–Thermal Interactions: Cooperation and Competition. IEEE Transactions on Haptics 18, 456–469 (2025).

62. P. J. Pardo, M. I. Suero, Á. L. Pérez, Correlation between perception of color, shadows, and surface textures and the realism of a scene in virtual reality. Journal of the Optical Society of America A 35, B130–B135 (2018).

63. M. Niu, C.-H. Lo, Do we see rendered surface materials differently in virtual reality? A psychophysics-based investigation. Virtual Reality 1–15 (2022). 10.1007/s10055-021-00613-3.

